# Comparative analysis of the Illumina Mouse Methylation BeadChip and Reduced Representation Bisulphite Sequencing for routine DNA methylation analysis of murine samples

**DOI:** 10.1101/2022.03.07.483250

**Authors:** Lochlan J Fennell, Gunter Hartel, Diane M McKeone, Catherine E Bond, Alexandra Kane, Barbara A Leggett, Ann-Marie Patch, Vicki LJ Whitehall

## Abstract

**Background:** Researching the murine epigenome in disease models has been hampered by the lack of an appropriate and cost-effective DNA methylation array. Until recently, investigators have been limited to the relatively expensive and analysis intensive bisulphite sequencing methods. Here, we performed a comprehensive, comparative analysis between the new Mouse Methylation BeadChip (MMB) and reduced representation bisulphite sequencing (RRBS) in two murine models of colorectal carcinogenesis, providing insight into the utility to each platforms in a real world environment.

**Results:** We captured 1.47×10^6^ CpGs by RRBS and 2.64×10^5^ CpGs by MMB, mapping to 13,778 and 13,365 CpG islands, respectively. RRBS captured significantly more CpGs per island (median 41 for RRBS versus 2 for MMB). We found that 64.4% of intra-island CpG methylation variability can be captured by measuring approximately one quarter of CpG island (CGI) CpGs. MMB was more precise in measuring DNA methylation, especially at sites that had low RRBS coverage. This impacted differential methylation analysis, with more statistically significantly differentially methylated CpG sites identified by MMB in all experimental conditions, however the difference was minute when appropriate thresholding for the magnitude of methylation change (0.2 beta value difference) was applied, providing confidence that both techniques can identify similar differential DNA methylation. Gene ontology enrichment analysis of differentially hypermethylated gene promoters identified similar biological processes and pathways by both RRBS and MMB across two murine model systems.

**Conclusion:** MMB is an effective tool for profiling the murine methylome that performs comparably to RRBS, identifying similar differentially methylated pathways. Although MMB captures a similar proportion of CpG islands, it does so with fewer CpGs per island. We show that subsampling informative CpGs from CpG islands is an appropriate strategy to capture whole island variation. Choice of technology is experiment dependent and will be predicated on the underlying biology being probed.

## Introduction

DNA methylation is the covalent modification of DNA to include a methyl group (CH3). This is a common epigenetic alteration that can govern chromatin accessibility, transcription factor activity, gene regulation and transcript expression. DNA methylation is deposited by DNA methyltransferase enzymes (DNMT), and removed by Ten-Eleven-Twelve enzymes (TET). Most DNA methylation occurs in the context of CG dinucleotides (CpG – cytosine-phosphate-guanine), with the cytosine nucleotide becoming methylated. Deregulation of the DNA methylation landscape is linked to several diseases. The CpG island methylator phenotype, which describes the widespread accumulation of DNA methylation at CpG islands, occurs in several forms of cancer, including colorectal and gastric cancers, and glioma (Fennell et al. 2019; Zouridis et al. 2012; Noushmehr et al. 2010; Liu et al. 2019). DNA methylation dysregulation also occurs in heart disease (Navas-Acien et al. 2021; Serra-Juhé et al. 2015), Alzheimer’s disease (Levine et al. 2015; Mastroeni et al. 2010), and rheumatoid arthritis (Nakano et al. 2013).

These associations have been borne out by studies utilizing the increasingly accessible of genome-wide DNA methylation technologies. In the late 1990s and early 2000s, PCR based methodologies were the primary modality of choice for investigating DNA methylation alterations. These approaches are limited in throughput, and thus stymie discovery-based investigations. Higher throughput In 2006, Illumina released the GoldenGate BeadArray Methylation assay, which was a microarray based approach that allowed for the simultaneous assessment of the DNA methylation state of ~1500 CpG sites that were located in the proximal promoters of ~370 genes (Bibikova and Fan 2009). The BeadArray platform has evolved over the past 15 years to include >850,000 CpG sites in its latest rendition, the EPIC array (Pidsley et al. 2016). The EPIC array, together with it’s predecessor, the 450K array, have been used extensively to characterize the DNA methylation landscape of normal and pathological states. This has led to several paradigm-shifting publications (Wockner et al. 2014; Hinoue et al. 2012; Horvath 2013; Nazor et al. 2012). These technologies are limited to human samples, and thus experimental biologists seeking to understand the mechanisms behind these associations using animal models must resort to other methodologies.

Transgenic murine models are an important tool for understanding the effects of certain genes on DNA methylation. A typical experiment might include knocking out a gene of interest, observing a phenotype and assessing the underlying DNA methylation profile of a given tissue. To achieve the latter, sequencing-based approaches such as whole genome bisulphite sequencing (WGBS) (Lister et al. 2009) or reduced representation bisulphite sequencing (RRBS) are usually employed (Gu et al. 2011). WGBS, if sequenced with sufficient depth, will cover >99% of CpGs in the murine methylome, but currently costs several thousands of dollars per sample. RRBS involves an enzymatic digestion at by MspI, which cuts DNA at CCGG motifs (Gu et al. 2011), and thus generates smaller fragments that map to CpG dense regions of the genome, reducing sequencing library size and ultimately sequencing costs at the expense of coverage of CpGs in other regions (Gu et al. 2011). The inclusion of an enzyme digest and size selection step also creates technical variability in the specific CpGs captured in each library preparation.

Illumina have recently developed a mouse DNA methylation microarray based on the BeadArray technology. This array captures >285,000 CpG sites. These sites are curated to include proximal promoter regions, and other regulatory regions of the murine methylome. In this study, we generate DNA methylation data from transgenic *Kras* and *Braf* mutant animals using reduced representation bisulphite sequencing (RRBS) and the Mouse Methylation BeadChip (MMB). In human colorectal cancers, both *BRAF* and *KRAS* mutation associate with DNA methylation dysregulation (Fennell et al. 2019; Hinoue et al. 2012; Weisenberger et al. 2006). As such, transgenic murine models recapitulating these mutations are ideal models for examining DNA methylation profiling technologies. We have performed a comprehensive comparative analysis to critically appraise this new platform and provide insights into the benefits and limitations of each technology.

## Methods

### Mouse models, sample collection and nucleic acid extractions

The *Braf*^*V637*^ and *KRAS*^*G12D-LSL*^ murine models were used for this study. The *Braf*^*V637*^ model is a cre-recombinase dependent conditionally activated model that facilitates the expression of the oncogenic *Braf* V637E allele. The V637E allele is analogous to the human V600E mutation. The *KRAS*^*G12D-LSL*^ conditionally activated model similarly expresses the oncogenic allele upon exposure to cre-recombinase. Both models were independently crossed with animals bearing the Villin-Cre^ERT2^ transgene. Villin-Cre^ERT2^ animals express the Cre^ERT2^ fusion gene under the guise of the Villin promoter, which is only active in the lower gastrointestinal tract. A single IP injection (75mg/kg) of tamoxifen facilitates the translocation of Cre^ERT2^ to the nucleus, where it induces recombination of the targeted alleles. We injected animals with tamoxifen at weaning and sacrificed them at two, eight and fourteen months of age. We also sacrificed age-matched littermates as wild type controls. Table 1 is a description of the animals used throughout the study. All animal experiments were approved by the QIMR Berghofer Animal Ethics Committee (P2178).

**Table 1:** Samples investigated in the study

| Sample Group | Age (months) | n (MMB) | n (RRBS) |
| --- | --- | --- | --- |
| Wild-type | 2 | 6 | 7 |
| Wild-type | 8 | 6 | 8 |
| Wild-type | 14 | 6 | 7 |
| *Kras* mutant | 2 | 3 | 4 |
| *Kras* mutant | 8 | 3 | 4 |
| *Kras* mutant | 14 | 3 | 4 |
| *Braf* mutant | 2 | 3 | 4 |
| *Braf* mutant | 8 | 3 | 3 |
| *Braf* mutant | 14 | 3 | 3 |

Tissue from the proximal small intestine was dissected at necropsy and cryopreserved in liquid nitrogen. DNA and RNA was extracted from fresh-frozen tissue samples using the AllPrep DNA/RNA/Protein minikit (Qiagen, USA) as per the manufacturer’s instructions. The same DNA extraction was used for both RRBS and MMB.

### Reduced Representation Bisulphite Sequencing and data processing

We generated single base resolution DNA methylation data using the Ovation RRBS Methyl-Seq System 1-16 (Tecan Life Sciences) with 100ng of input DNA. This approach employs the *Msp*I restriction enzyme to digest DNA at the CCGG motif, which is highly enriched in CpG dense regions of the genome. This generates a library of small (>300bp) fragments that are rich in CpG content. Sequencing libraries are generated from these fragments, bisulphite converted and sequenced. For this study, we sequenced these libraries to a target depth of 30 million single end 100bp reads per sample on an Illumina NovaSeq instrument.

Data were processed as per Fennell et al (2021). Briefly, sequencing reads were inspected for quality using FastQC (v0.11.7). Reads were trimmed to remove sequencing adaptors and poor quality bases using TrimGalore (v0.6.6, (Krueger et al. 2021)). Reads were then aligned to the murine methylome (mm10) using Bismark (v0.20.0) and methylation calculated from alignments using the methylation_extractor and bismark2bedGraph functions of Bismark (v0.20.0, (Krueger and Andrews 2011)). Data were imported into the R environment using the methylKit package (v1.14.2, (Akalin et al. 2012)). Using methylKit, CpG sites covered by < 10 reads in any sample were discarded to generate a consensus dataset. We also performed analyses at various levels of coverage to determine how many CpGs are lost when coverage requirements increase.

### Mouse Methylation BeadChip

Genome-wide DNA methylation was performed using the Illumina Infinium Mouse Methylation BeadChip (Illumina, San Diego, CA, USA) following the standard manufacturer’s protocol. 500ng of high-quality genomic DNA was bisulfite converted using the Zymo EZ-96 DNA Methylation kit (Zymo Research, Irine, CA, USA). Bisulfite converted samples were then amplified, fragmented, purified and hybridized onto the Mouse Methylation BeadChip according to the manufacturer’s standard protocol. The arrays were washed and scanned using the Illumina iScan System. Mouse Methylation BeadChips were processed at Australian Genome Research Facility (AGRF), Melbourne For analysis, idat files were imported into R using EnMix (v1.25.1, (Xu et al. 2016)). Data was normalized using BMIQ and filtered by detection P and for sex chromosomes. For detection P filtering, we masked values that had a detection P > 0.05, and removed the probe entirely if >50% of samples had a detection P > 0.05.

### Data Annotations

To annotate CpGs with respect to whether they reside in CpG islands, we downloaded the mm10 cpgIslandExt table from the UCSC table browser. The cpgIslandExt table contains annotations of CpG islands, where a genomic region is a CpG island if it meets the following criteria: having >50% GC content, a length of >200bp and a ratio of observed to expected CG dinucleotides of >0.6. We assigned each CpG island a unique identifier, and examined overlaps between CpGs on MMB and RRBS using the intersect function of bedtools (v2.29.0, (Quinlan and Hall 2010)).

Annotations of candidate regulatory elements were downloaded from the murine ENCODE project web portal (Mouse ENCODE Consortium et al. 2012). These annotation tracks were generated by the ENCODE consortia through evaluation of ChIP-Seq data pertaining to histone modifications, and DNase-hypersensitivity sequencing. These annotations included promoter-like (PLS; within 200bp of TSS, ++ DNase accessibility and H3K4me3 signal), proximal enhancer-like (pELS; within 2000bp of TSS, ++ DNase accessibility and H3K27ac signal, low H3K4me3 if <200bp to TSS), distal enhancer-like (dELS; >2000bp from TSS, ++ DNase accessibility and H3K27ac signal), CTCF-only (high DNase and CTCF, low H3K27ac and H3K4me3) and DNase-H3K4me3 (>200bp from TSS, ++ DNase accessibility and H3K4me3 signal). We examined overlaps between CpGs and candidate regulatory elements using the intersect function of bedtools (v2.29.0).

Gene promoter annotations were generated using the annotatePeaks function of the HOMER tool (v4.8, (Heinz et al. 2010)) and the mm10 reference genome. CpGs that were subsequently annotated as promoter and ascribed to a gene were used for downstream gene promoter analysis.

### Statistical analyses

To identify a subsets of similarly methylated CpGs in each CpG island we used the *Cluster Variables* procedure in JMP (v16, SAS Institute, Care NC, USA). The algorithm starts with a set of variables and splits them into clusters comprising highly correlated variables, and identifies a single representative variable which explains the largest proportion of the variance in the cluster. To evaluate the variability of DNA methylation at single CpG sites within experimental groups we used the F-test method as employed in the matrixStats R Package (v0.61.1). This method tests whether the variances across two groups are equal. Data is presented as the log(variance) ratio, with <0 being representing less within group variability on MMB and >0 representing greater within group variability on MMB.

To assess the degree of correlation between MMB and RRBS we first generated a consensus dataset that contained DNA methylation measurements of the same CpG on both platforms. For each sample, we then performed linear regression analysis on the entire consensus data set, comparing measurements on MMB with measurements on RRBS. For differential methylation analysis we used the Limma R package (v3.14, (Ritchie et al. 2015)). For gene ontology enrichment analysis we used the ClusterProfiler R package (v3.16.1, (Wu et al. 2021)). In comparing the number of differentially methylated CpGs or gene ontology terms, we the X^2^ method with Yates correction.

## Results

### Coverage of MMB and RRBS

We first sought to examine the coverage of both platforms. After filtering, 264,145 CpGs were captured by the MMB, of which 207,468 were captured in all samples (no sample level filtering). For RRBS, at our threshold of 10X coverage, we captured 1,487,242 individual CpG sites. We also examined the number of CpGs covered at sequencing depths from 5X to 40X (Figure 1). As expected, there was a decrease in the number of CpGs captured with increasing sequencing depth filtering. These data indicate that the depth to which the library is sequenced is an important factor in RRBS experimental design.

**Figure 1:**
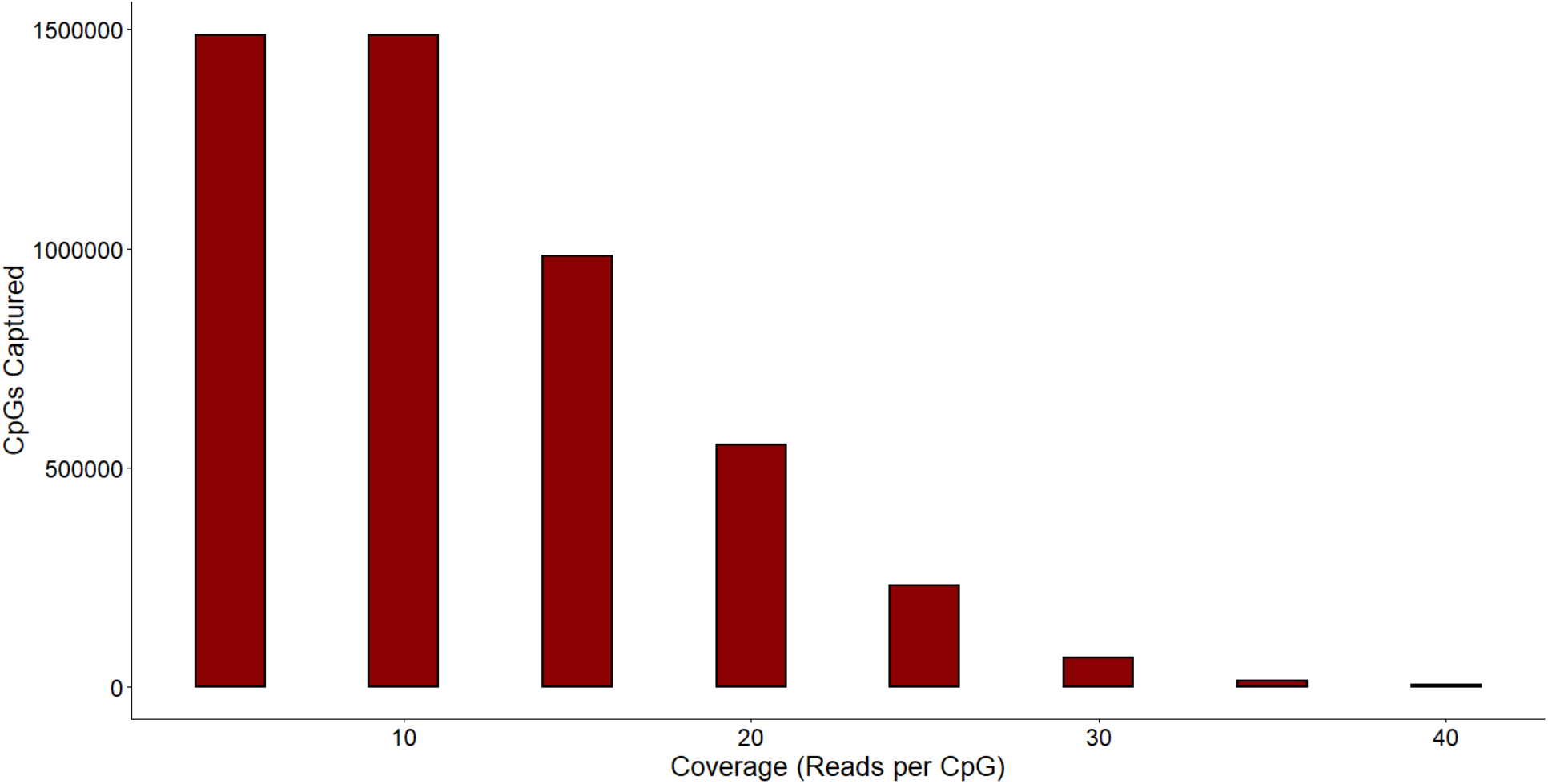
CpGs captured by reduced representation bisulphite sequencing at depths ranging from 5X to 40X. Data represents the number of CpGs captured at each depth, in all samples.

Next, we examined the coverage of CpG islands (CGI) by each technology. In this study, we adopted the UCSC definition of a CpG island, which describes islands as: having >50% GC content, a length of >200bp and a ratio of observed to expected CG dinucleotides of >0.6. Using this criteria, we identify 17,017 CpG islands in the murine genome. RRBS covered 13,778 CpG islands (~80% of all annotated islands) with at least one CpG site. The median number of CpG sites covered per CpG island was 41. MMB covered a similar number of CpG islands (13365), however median number of CpGs per CpG island on the array was only 2. CpGs covered in RRBS were enriched for CpG islands compared with those on the MMB, with 48.9% of RRBS CpGs residing in CpG islands, versus 11.5% on the MMB. These data indicate that the breadth of the array, at the level of CpG islands, is similar to RRBS, however the intra-island coverage is substantially lower.

To examine the breadth of coverage of MMB and RRBS in regulatory regions we downloaded candidate cis-regulatory elements from the murine ENCODE project. These elements are subdivided into promoter-like, proximal enhancer-like, distal enhancer-like, CTCF-only and DNase-H3K4me3. The murine ENCODE project describes 23,762 PLS, 72,964 pELS, 209040 dELS, 23836 CTCF-only, and 10,383 DNase-H3K4me3 regions.

MMB has probes that cover 2.4% of PLS (571 regions), 6.8% of dELS (4927 regions), 0.73% of pELS (1542 regions), 0.88% of DNase-H3K4me3 regions (211 regions), and 5.12% of CTCF-Only sites (532 regions). In total, MMB has probes that cover 7,783 independent candidate cis-regulatory regions.

By comparison, at 10X coverage, the RRBS consensus dataset contained CpGs that mapped to 57,068 unique candidate cis-regulatory elements, including 60.9% of PLS (14,480 PLS regions), 29.4% of pELS (21,450 regions), and 8.8% of dELS (18356 regions).

### Appropriate subsampling of CpGs within CGIs can capture most intra-CpG island variability

RRBS captures significantly more CpG sites per CGI. We sought to determine whether capturing these CpG sites yielded additional insight into the DNA methylation profile of the CGI, or whether they were methylated similarly and provided redundant information. To better understand the variability within CGI we first calculated the variance of methylation across CGI in each sample, and then calculated the mean variance at each CGI across the entire dataset (Figure 2). The positive-skew in the distribution of mean variances indicates that most CGIs are relatively stable, with little variability in DNA methylation between CpGs residing in the same island. Figure 2B-I depicts CGI with increasing intra-island variability. We hypothesized that CGI length and CGI length to number of CpG sites may influence intra-island variability, with longer CGIs and those represented by fewer CpGs being more variable. To test this, we regressed the variance of each island against the length (bp) and the CGI Length to number of CpG sites ratio (CpG:CGI-Length ratio). While both CGI length and CGI-Length:CpG ratio were highly significantly associated with increased intra-island variability (1.75×10^−6^ and 2×10^−16^), the R^2^ was low (R^2^ ≈ 0.01), indicating that most of the variability is ascribed to other factors.

**Figure 2:**
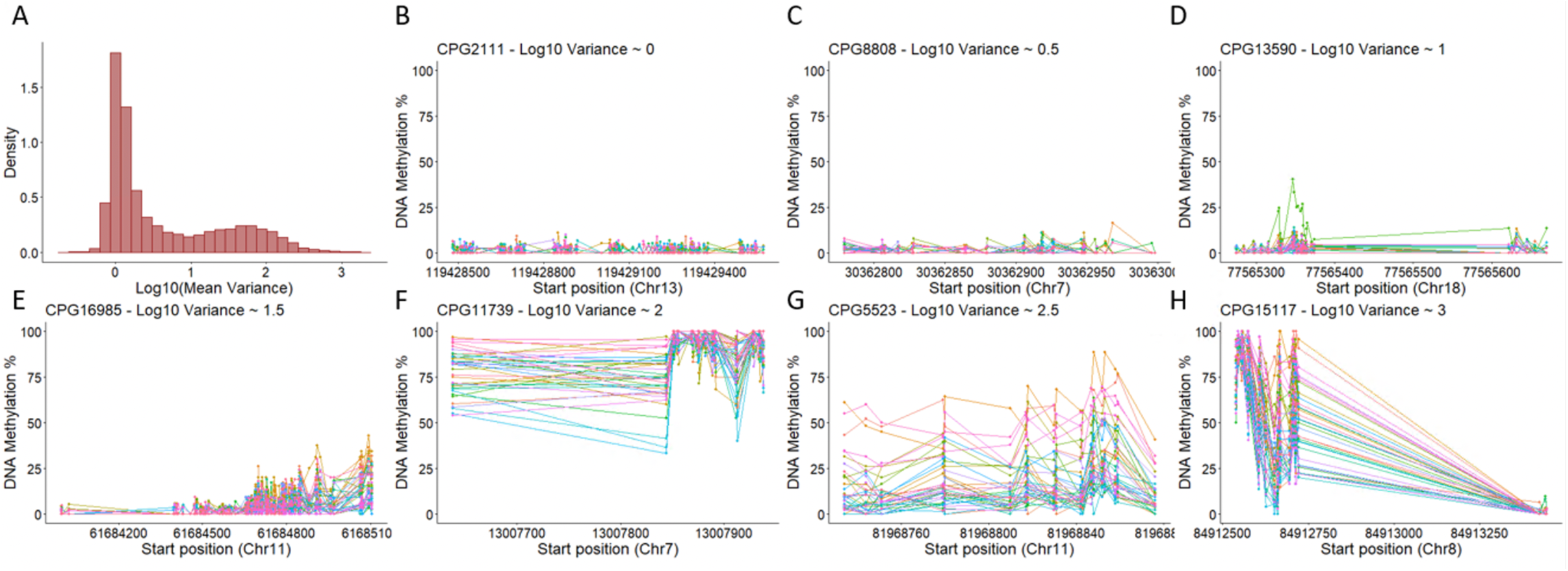
A: Variance of DNA methylation across CpG islands containing multiple CpG sites. The histogram represents the distribution of the log10(variance) between CpG sites in all CpG islands with >1 CpG. DNA methylation as measured by RRBS. Each dot and line represents methylation in an individual sample B-I: Representative DNA methylation profiles of CpG islands with low variance (B) through to CpG islands with extreme intra-island variability (H).

Next, we examined the effect of DNA methylation level on the stability of DNA methylation across the island. DNA methylation was most consistent across CGI when islands were near entirely methylated or demethylated (Figure 3). We quantified this statistically by calculating the absolute difference between average methylation across the island and hemimethylation (50%). As distance from hemimethylation increased (toward either 0 or 100% methylation) variability across the island significantly decreases (P=2×10^−16^, R^2^≈ 0.76).

**Figure 3:**
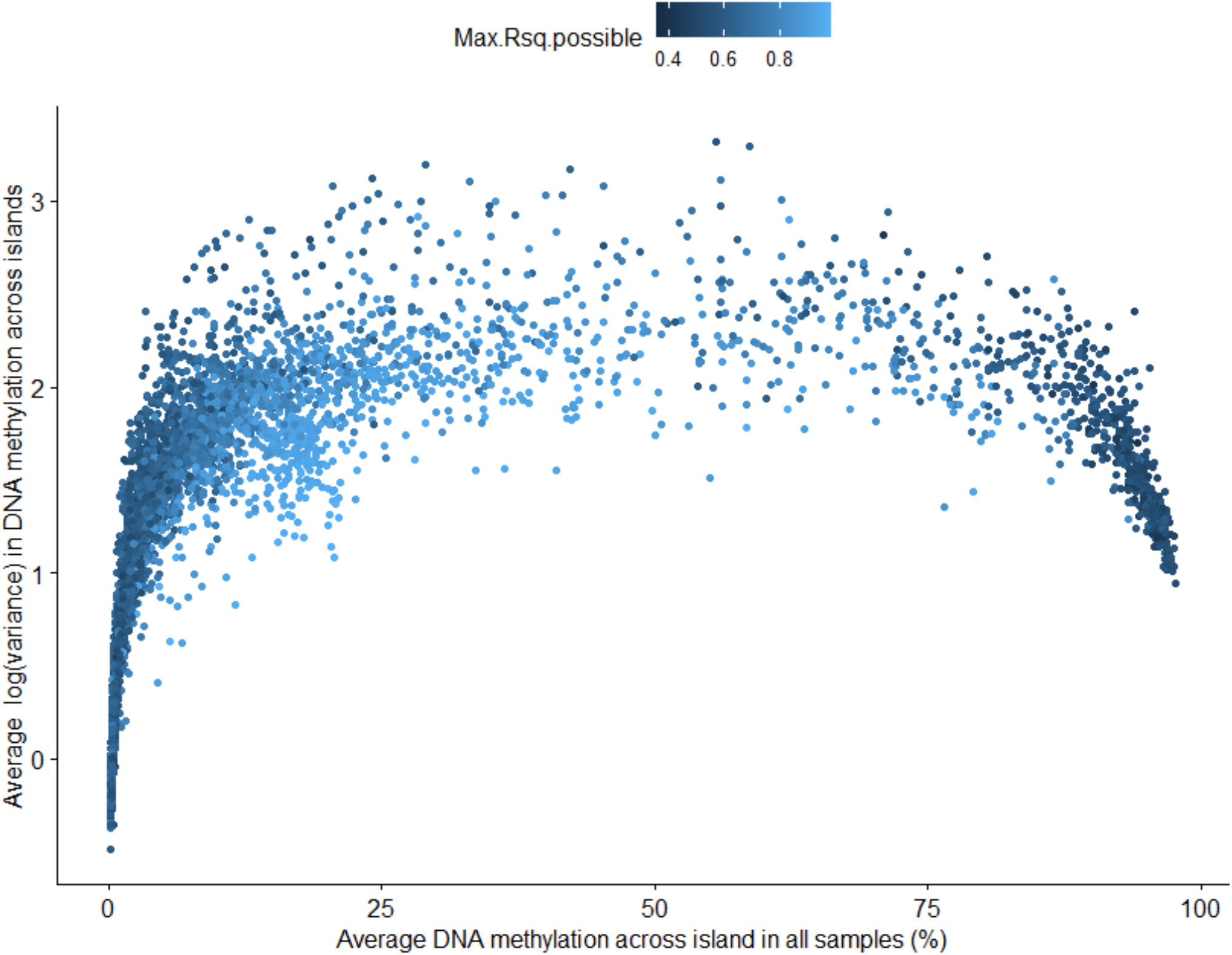
Average DNA methylation (%), as measured by RRBS, of each CpG island plotted against the intra-CpG island DNA methylation variability. Each dot represents a single CpG island. Dots are colored by the maximum r-sq achieved by variable clustering, where a high value indicates that the variability across the island can be well captured by a subset of CpGs.

To identify whether most of the variability between CpGs in an island could be captured by a smaller subset of CpGs we performed variable clustering. Variable clustering captures subsets of highly correlated variables, to which most of the variance within a cluster can be explained by a single variable. The average CGI in our RRBS dataset contained 52.59 CpG sites, and 13.4 variable clusters. We calculated how much of the total variability within a CGI could be explained by capturing only data from representative CpGs from variable clusters within an island. Using data from only the most representative CpGs of variable clusters within a CGI, we can recapitulate an average of 64.6% of the variability within a CpG island, and reduce the size of the total dataset pertaining to CGIs from 702,978 CpGs to 179,166 CpGs.

We observed that CGIs with low variability have lower % variance explained by variable clusters (Figure 3), and require a greater number of clusters to explain the variance across the CGI. This is likely due to the small variance, most of which is produced by noise rather than biological signal. In contrast, highly variable CpG islands, which reflect biological variability across the island, can be well explained by capturing several CpGs (Figure 3). These data indicate that highly variable CGIs can be accurately summarised by profiling a subset of carefully selected CpGs within the CGI.

### Base-resolution methylation profiling via MMB versus RRBS

Next, we sought to assess the correlation between methylation values attained at base-resolution using MMB and RRBS. After coverage filtering (>10 reads per CpG), our RRBS experiment covered 1.48×10^6^ CpG sites. 25,548 filter-passing CpGs were covered by both MMB and RRBS, representing 12.3% and 2.44% of CpGs covered by the MMB and RRBS, respectively. We extracted methylation values for these CpG sites from both platforms and computed sample-wise correlation analysis. In CpGs covered by both assays, we observed high concordance between both platforms (Pearsons R^2^ >0.95 in all samples, Figure 4).

**Figure 4:**
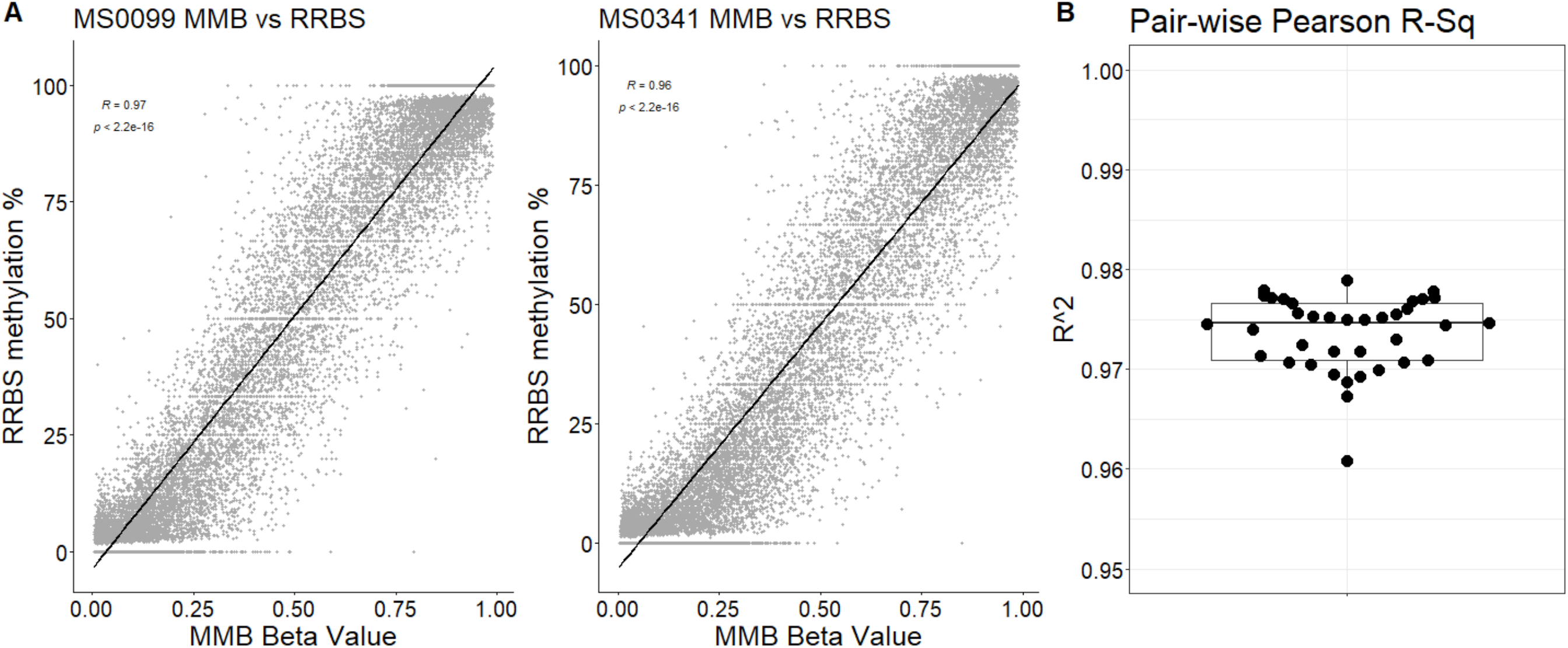
Regression analysis revealed significant genome-wide concordance of DNA methylation between MMB and RRBS. A: Representative DNA methylation measurements as captured by the MMB and RRBS in CpG sites that were common to both platforms. B: Pearsons’ R^2^ values for same sample whole methylome comparisons on RRBS and MMB

Next, we examined correlations between methylation values obtained on MMB and RRBS at individual CpG sites. There was substantial inter-CpG variability in the correlation between MMB and RRBS (Pearson’s R^2^ range: 0-0.98). We hypothesized that the correlation between readouts obtained on each platform may be weakest at measurement extremes (fully (un)methylated). To examine this hypothesis we calculated the average beta value obtained by MMB for each probe, and binned them in beta value increments of 0.05. We observed that correlations between MMB and RRBS followed a unimodal distribution centered around an average beta value of ~0.4 (Figure 5)

**Figure 5:**
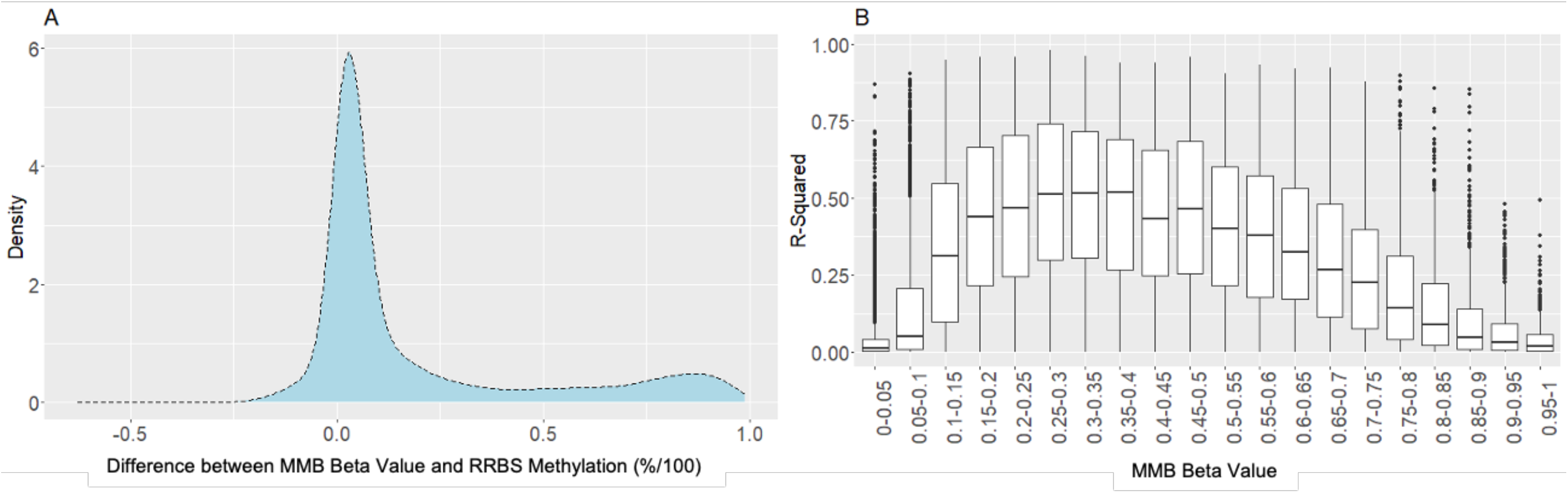
A: Distribution of the difference between the mean MMB beta value and the mean RRBS methylation % at individual CpG sites. B: Correlation of DNA methylation between MMB and RRBS (Rsq) versus the average beta value for the same CpG. CpGs were binned in increments of 0.05.

### Clustering and differential DNA methylation analysis of common probes

Most studies attempt to identify differential DNA methylation between experimental groups. We next sought to examine the ability of both technologies to identify epigenetic drift and differentially methylated CpGs. It is well established that methylation at specific loci shifts in accordance with advancing age (Fennell et al. 2021; Horvath 2013; Teschendorff et al. 2013). For epigenetic drift analyses, we regressed the beta values (or % methylated reads) against age (days) for CpGs that were assayed in both technologies. The Illumina Methylation array vastly outperformed RRBS in this respect, identifying statistically significant epigenetic drift in 3,274/25,396 overlapping CpGs (Figure 6). By contrast, RRBS identified a mere 278/25,396 CpGs. Furthermore, 92.4% of CpGs that were identified as changing with age by RRBS were also detected by MMB.

**Figure 6:**
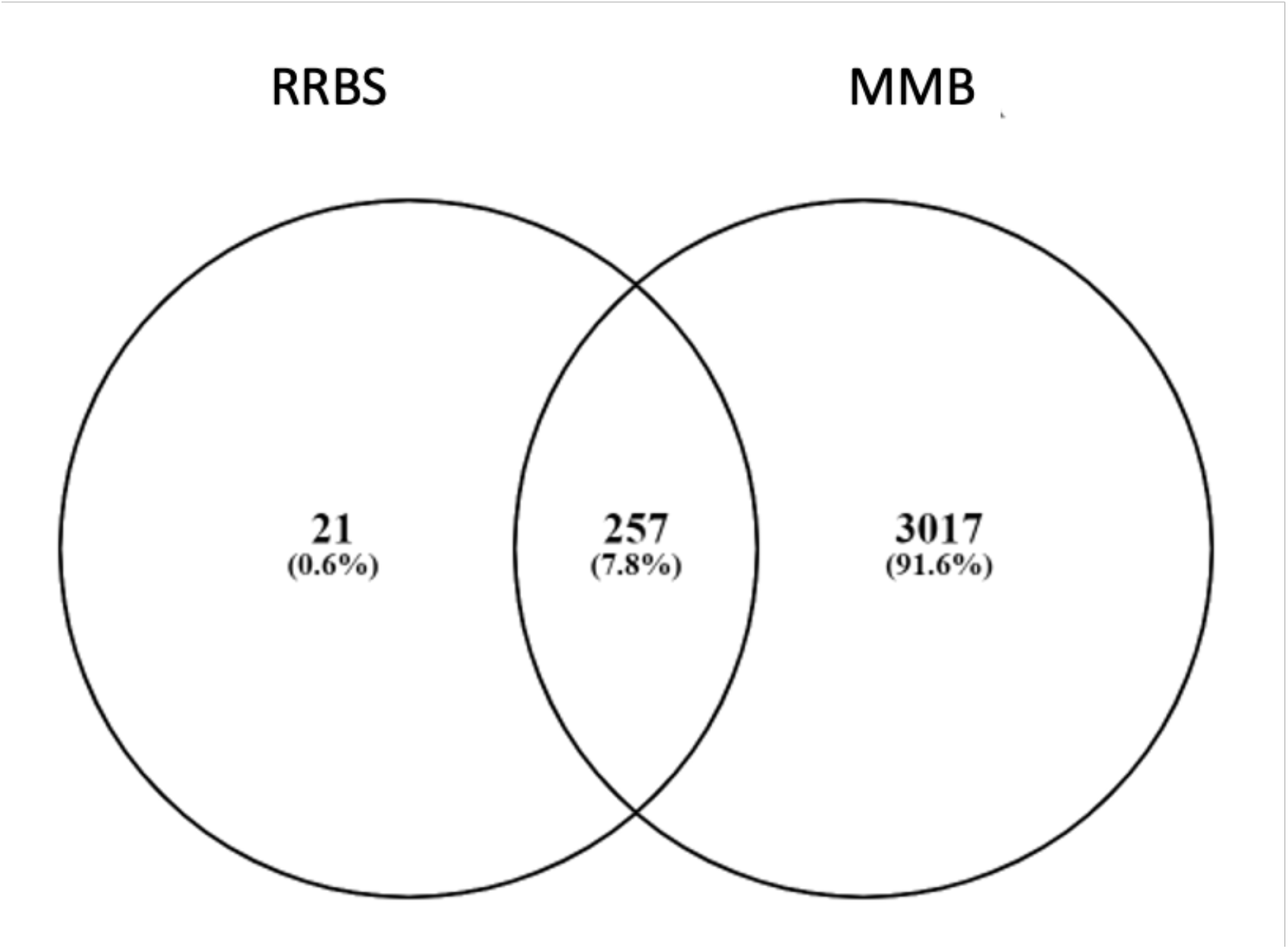
Overlap between genes common to both profiles and identified as significantly age-associated in wild-type samples by either RRBS, MMB or both.

We also performed differential DNA methylation analysis between *Kras or Braf* mutant animals and their wild-type counterparts at 8 and 14 months. In keeping with our epigenetic drift analysis, MMB detected significantly more statistically differentially methylated loci (Table 2). Most of the differences detected by MMB were small. By applying the common |Δ| 0.2 beta value threshold for calling differential methylation, the discrepancies in differential methylation calling on the two platforms shrank substantially (Table 2), indicating that the MMB may be more suited for detecting more subtle DNA methylation alterations.

**Table 2:**
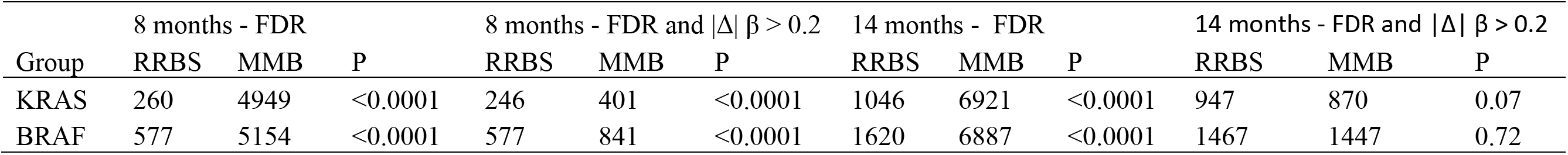
Number of differentially methylated CpG sites between 8 and 14 month Braf and Kras mutant animals versus wild type animals as assessed by RRBS and MMB in CpG sites that were common to both techniques.

### The Mouse Methylation BeadChip is more precise than RRBS

We hypothesized that the within-group variation in detected methylation may be smaller on the MMB platform compared to RRBS, thus improving our ability to detect more subtle age and genotype associated changes. To explore this we analysed within-group variance on each platform at single CpG resolution. For each experimental group and for most CpG sites (Figure 7), we observed substantially less variability between samples on the MMB. For both *Braf* and *Kras* mutant samples at 8 months, variability was significantly greater on the RRBS platform when compared with MMB, with 3,222 (*Braf* mutant animals) and 2,760 CpGs (*Kras* mutant animals), having a greater intra-group variability on RRBS when compared to MMB. By contrast, the RRBS was less variable than MMB at 19 and 20 CpGs, respectively. The same trend was observed at 14 months (Figure 7). These data support the hypothesis that RRBS generated DNA methylation data is more variable than MMB, and thus is limited in detecting smaller DNA methylation changes.

**Figure 7:**
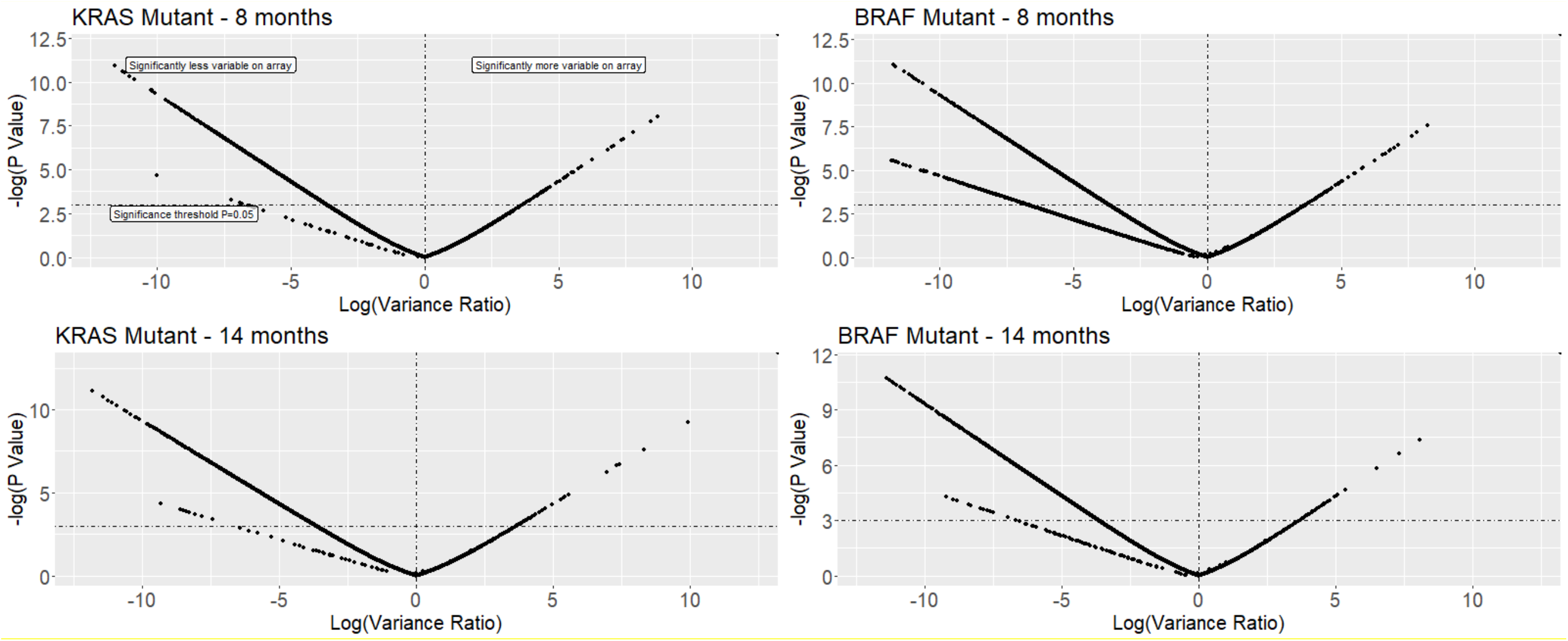
Log(Variance Ratio) of CpG sites common to both platforms by experimental group. CpGs with a log(Variance ratio) of <0 have lower within group variability on the MMB and >0 on RRBS. For all experimental groups, most CpGs had lower within-group variability on MMB. Log(Variance Ratio) calculated by F-Tests.

### Detection of common pathway alterations

Gene promoter and pathways analysis is commonly used to identify processes that are driving phenotypes of interest. We sought to assess whether each technology could independently identify differentially methylated promoters and pathways in our *Braf* and *Kras* mutant murine models. We performed differential methylation analysis on each platform, using the entire quality-filtered dataset (rather than just CpGs shared by each technology) in each case, to emulate the situation where the investigator has chosen a particular technology from the outset. In keeping with earlier analyses, we considered CpGs to be differentially methylated if the FDR was < 0.05 and there was an absolute difference in DNA methylation of 0.2 (MMB) or 20% (RRBS). Both technologies detected a large number of significant DNA methylation alterations in each condition versus wild-type animals. Unsurprisingly, due to the higher number of CpGs, RRBS detected more differentially methylated CpG sites than MMB in most comparisons (Table 3). The proportion of differential methylation when compared to the total number of CpGs captured by each technique was between 1 and 3%.

**Table 3:** Differential DNA methylation in Braf and Kras mutated intestine as detected by MMB and RRBS.

| Mutation | Direction | 8 months |  | 14 months |  |
| --- | --- | --- | --- | --- | --- |
|  |  | RRBS | MMB | RRBS | MMB |
| *Braf* | Hypermethylated | 11253<br>(0.76%) | 4423<br>(1.7%) | 46757<br>(3.14%) | 6276<br>(2.41%) |
|  | Hypomethylated | 13908<br>(0.93%) | 9423<br>(3.61%) | 16399<br>(1.1%) | 7808<br>(2.99%) |
| *Kras* | Hypermethylated | 5288<br>(0.36%) | 1962<br>(0.75%) | 30625<br>(2.06%) | 3729<br>(1.43%) |
|  | Hypomethylated | 2127<br>(0.14%) | 2263<br>(0.87%) | 5597<br>(0.38%) | 3312<br>(1.27%) |

<sup>1</sup>The entire probeset was included in the analysis of the MMB and all CpGs with $>10X$ coverage for RRBS. Data represents the number of differentially methylated CpGs and the proportion of differentially methylated CpGs when compared to all CpGs covered.

As these models represent intestinal oncogenes that are usually associated with the CpG island methylator phenotype, we next segregated the data by whether the CpG resided in a CGI or not. As outlined in earlier paragraphs, 727,509 (48.9%) and 30,385 (11.6%) CpGs reside in CGIs in our RRBS and MMB datasets, respectively. On both platforms, we observed the expected pattern of distribution of hyper(hypo)methylation events (Figure 8), with hypermethylation concentrated in CpG islands and hypomethylation in non-island associated genomic regions.

**Figure 8:**
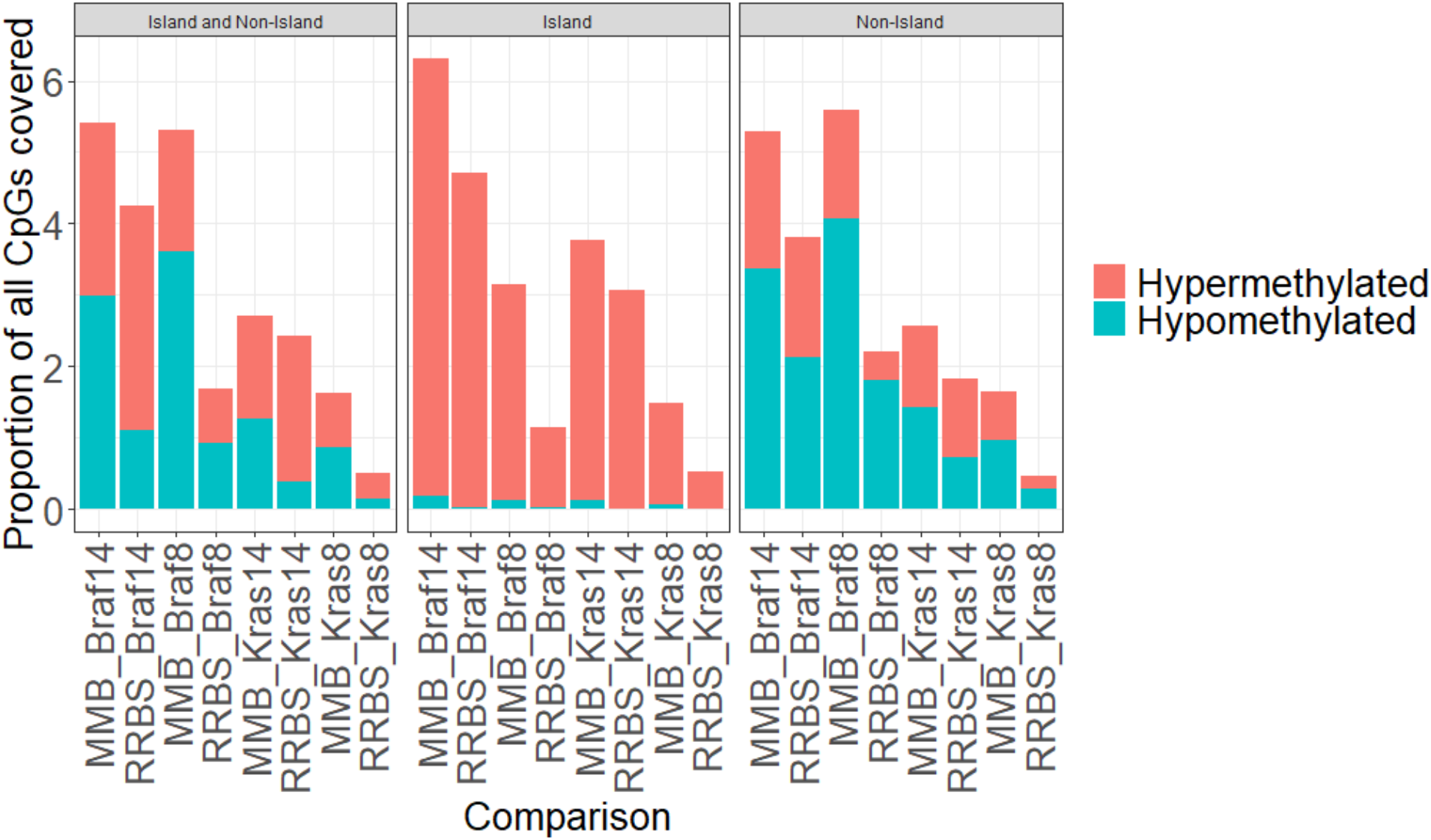
Proportion of all CpGs covered that were differentially methylated on each platform, for each group. Differential methylation is faceted by whether the CpG resides in an island or not.

Next, we performed gene ontology analysis on hypermethylated promoters in 14 month *Braf and Kras* mutant mice on MMB and RRBS. For each comparison we observed an enrichment for hypermethylation at promoters encoding genes involved in differentiation and development (Figure 9), consistent with our prior work (ref fennell et al). We observed substantial overlap in the gene ontology processes identified by MMB and RRBS) (64.4% and 51% for *Braf* and *Kras*, respectively), and ~65% of the top 50 enriched terms for both models were identified by both RRBS and MMB (Figure 9). These data indicate that both technologies can reliably detect differentially methylated pathways that are relevant to the disease model.

**Figure 9:**
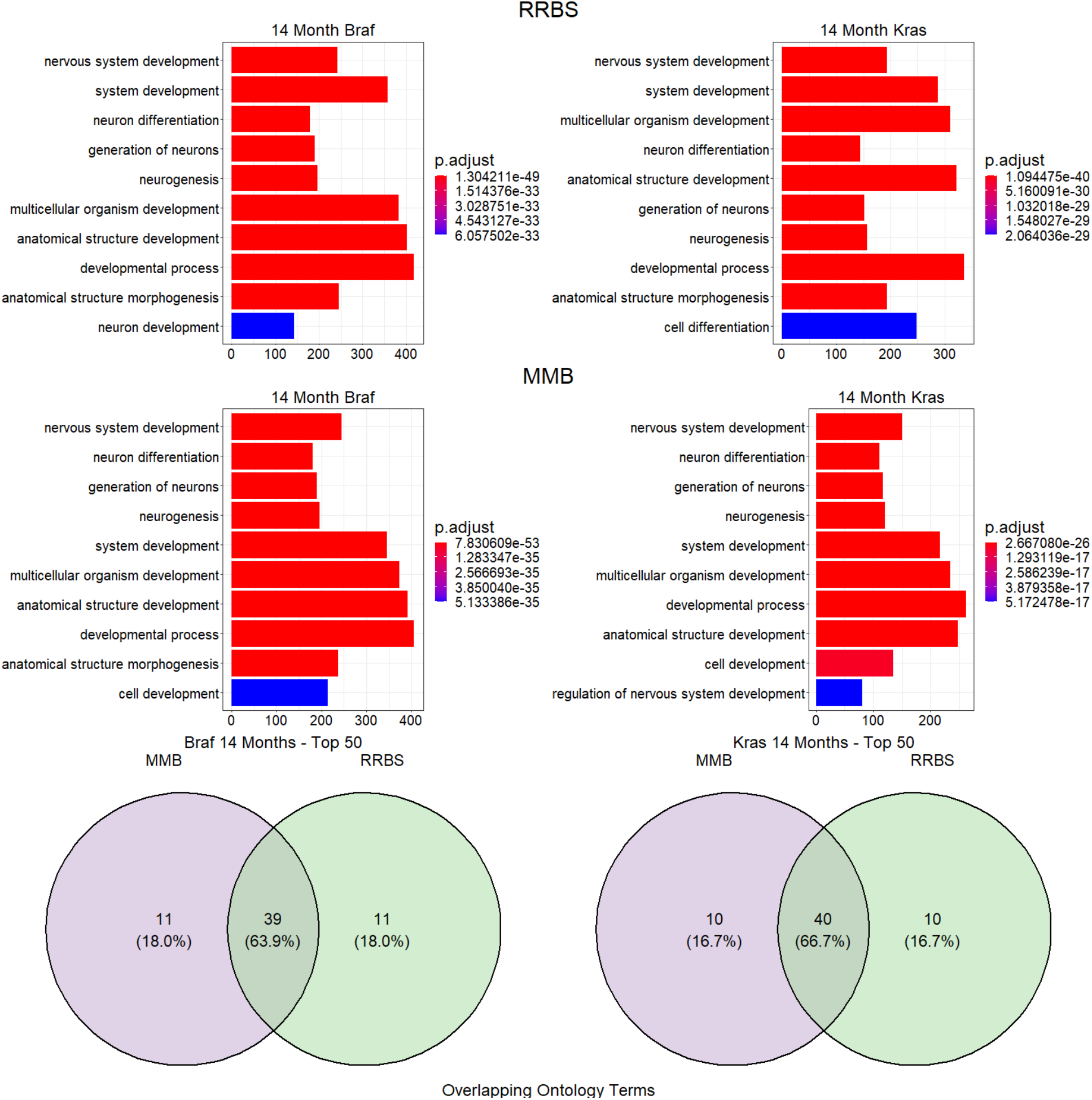
Top: Gene ontology enrichment analysis for differentially hypermethylated promoters detected in Braf or Kras mutant 14 month old mice versus wild-type littermates as captured by RRBS and by MMB. Bottom: Overlap of gene ontology terms determined by analysis of MMB and RRBS data.

## Discussion

DNA methylation is an important epigenetic modification that can govern chromatin state and gene transcriptional programs. Several diseases alter the DNA methylation landscape (Robertson 2005), and understanding the basis for these modifications is important for understanding disease etiology. Murine models are ideal for investigating disease, however genome-scale DNA methylation analyses have been limited to bisulphite-based sequencing approaches, such as whole-genome bisulphite sequencing, which is prohibitively expensive, and reduced-representation bisulphite sequencing, which has a coverage bias for CpG island associated CpGs. Here, we have compared a new array-based DNA methylation analysis tool, the Mouse Methylation BeadChip, to reduced representation bisulphite sequencing. We report that MMB performs comparably to RRBS. MMB produces precise measurements of DNA methylation and is able to detect disease specific DNA methylation alterations.

Both techniques capture a similar number of CpG islands, however the Mouse Methylation BeadChip has significantly fewer CpGs per island. There is some evidence that suggests there is a high level of correlation between the DNA methylation of CpG sites within a CpG island, especially when the CGI is small (<400bp) (Zhang et al. 2015). However, most of this research is confined to humans, and it is not clear whether this was also true in murine models. Here, we have sought to assess whether increased intra-CpG island coverage by RRBS might contain biologically relevant DNA methylation, or whether additional CpG coverage within an island is redundant, with DNA methylation of a few CpGs being sufficient to infer DNA methylation of the entire island. This is an important distinction, given that the CpG islands are typically only represented by one or two probes on the MMB. Here, we report that most CpG islands had relatively low intra-island methylation variability. As the mean methylation of a CpG island approached 50% we observe significant variability in DNA methylation between CpGs within the island, and thus it is important to consider the validity of generalizing MMB data generated from a small number of probes to the entire CpG island at intermediate levels of DNA methylation.

In sequencing experiments, investigators are often faced with the coverage versus cost trade-off, whereby increased coverage and sequencing depth is associated with read-outs that are more reliable but can dramatically increase the overall cost of the experiment. In this study, we considered 10X coverage to be an appropriate cut-off for calling DNA methylation fractions, allowing us to profile ~1.5 million CpG sites. We note that as we increase our coverage stringency, we obtain more accurate methylation fractions that converge on the methylation read-out obtained for the same sites on MMB. If the CpG site(s) of interest are covered by MMB, this technology may offer a more precise read-out that is more cost effective than deep sequencing of bisulphite sequencing libraries. We also note that as little as 100ng is sufficient for generating RRBS libraries. In contrast MMB requires >250ng. Investigators should keep these factors in mind when considering which technology to use. Ultimately, the underlying biological processes being probed and the resources of the laboratory will guide these decisions.

One common application is differential methylation analysis. Here, the question is whether there is a statistically significant difference in DNA methylation at a given site between two experimental groups. Small, consistent differences can often yield significant p values, but are have little biological relevance. We and others also apply a Δ methylation cut-off, usually of 0.2 for methylation arrays or 20% for methylation sequencing. One of our key findings here is that although MMB can detect substantially more differential methylation between experimental groups at a purely statistical level, this difference is negligible when we apply an absolute methylation change cut-off. For most animals experiments, investigators will be far more interested in large changes in DNA methylation at a given loci, thus these data should provide confidence that both methods, if covering the loci of interest, will detect these alterations. For more niche applications where subtle changes in DNA methylation are of interest (for example, when profiling a locus that is methylated in a subpopulation of cells in bulk samples), MMB may be more appropriate.

The expertise and availability of bioinformatics infrastructure and support should be considered when choosing an approach. RRBS is a sequencing based approach, and therefore will generate millions of sequencing reads that must be preprocessed, aligned and methylation-called. This process is computationally intensive, and requires specialized bioinfomatic expertise. MMB data can be processed entirely in the R environment, with several well described R packages facilitating array processing (Xu et al. 2016; Aryee et al. 2014; Müller et al. 2019). By contrast, RRBS data, as with all large sequencing experiments, must be first processed using command-line tools, presenting an additional technical barrier for inexperienced users. It is not possible to process these data on standard computers, and requires access to compute infrastructure. MMB generates a comparably smaller amount of data, and can be processed on high-end desktop computers. Basic array analysis and processing is more accessible to non-bioinformatically trained scientists, and can also be performed on platforms with graphical interfaces (ie. Galaxy). The availability of specialized expertise and infrastructure is an important element that should be considered when choosing a technology.

In our final analysis, we performed a side-by-side differential methylation and pathways analysis on both platforms to confirm that similar biological signatures would emerge with both techniques. We have previously shown that *Braf* mutation in humans and in mice results in widespread accumulation of DNA methylation at CpG islands (Fennell et al. 2021, 2019; Bond et al. 2018). We have also shown that *Kras* mutation can generate a similar but distinct methylator phenotype. Using RRBS and MMB we have recapitulated these signatures, validating both the animal models and the approaches used to assess DNA methylation.

Here we have comprehensively evaluated the performance of the Illumina Mouse Methylation Array in comparison to Reduced Representation Bisulphite Sequencing in two models of murine colorectal carcinogenesis using matched samples. In CpG sites that are covered by both platforms, we report high correlations between DNA methylation profiles. IMMA could detect a greater number of statistically differentially methylated sites than RRBS; IMMA was able to detect smaller methylation alterations due to the increased precision of DNA methylation readouts, especially in regions that are poorly covered by RRBS. However, this difference was not significant when we imposed a minimum methylation change threshold, as is common in the literature. To conclude, IMMA is a valuable addition to the toolbox of experimental epigeneticists that performs comparably to sequencing based methodologies. IMMA is highly suited to applications requiring highly precise methylation calls, and the detection of small methylation differences between experimental groups.

